# Early-life environment programs reproductive strategies through epigenetic regulation of *SRD5A1*

**DOI:** 10.1101/2020.09.16.299560

**Authors:** Ben Bar-Sadeh, Or Eden, Lilach Pnueli, Kurshida Begum, Gregory Leeman, Richard D. Emes, Reinhard Stöger, Gillian R. Bentley, Philippa Melamed

## Abstract

Reproductive function and duration of the reproductive life span are phenotypically plastic and programmed in response to the early-life environment. Such adaptive responses are described and rationalized in life history theory in the context of resource availability, but the molecular mechanisms responsible have remained enigmatic. In this study, we hypothesized that epigenetic modifications underlie adaptive reproductive strategies, and found distinct methylation patterns in buccal DNA of Bangladeshi women who grew up in Bangladesh or the UK. The later pubertal onset and lower ovarian reserve associated with Bangladeshi childhood was seen to correlate with more numerous childhood infections, so we adopted a mouse model of pre-pubertal colitis to mimic these conditions. These mice have a similarly-altered reproductive phenotype, which enabled us to determine its mechanistic basis. Several genes encoding proteins with known functions in follicle recruitment were differentially expressed in the mice ovaries, and were also differentially methylated in the women’s buccal DNA. One of these, *SRD5A1* which encodes the steroidogenic enzyme 5α reductase-1, was down-regulated in the mice ovaries and hyper methylated at the same putative transcriptional enhancer as in the women’s DNA; the levels of methylation correlating with gene expression levels. *Srd5a1* expression was down-regulated also in the hypothalamus where 5α reductase-1 catalyzes production of neurosteroids that regulate gonadotropin releasing hormone (GnRH). Chemical inhibition of this enzyme affected both GnRH synthesis and release, and resulted in delayed pubertal onset *in vivo*. The activity of 5α reductase-1 in hypothalamus and ovary and the sensitivity of *SRD5A1* to epigenetic regulation attest to its role in directing long-term physiological strategies in response to environmental conditions. In the reproductive axis, this includes timing of pubertal onset, adult reproductive function and duration of the reproductive lifespan.

## Introduction

Reproductive function is plastic, responding and adapting to environmental signals with changes in age of sexual maturation, hormone levels, rates of ovulation and fertility, as well as length of a woman’s reproductive lifespan^1,2^. The adult reproductive phenotype is particularly sensitive to the early life environment and is largely programmed by mid-childhood. This was evident in studies on Bangladeshi migrants, in which we found that women who spent their childhood in Bangladesh, at least until aged 8 years, experienced later pubertal onset, earlier menopause and had a lower ovarian reserve than Bangladeshi women who grew up in the UK^3–6^. Children who migrated at a younger age, or second-generation Bangladeshis in the UK, had similar reproductive phenotypes to their European ethnic neighbors, while women who were adults at migration maintained the “Bangladeshi childhood” reproductive phenotype even after many years in the UK. Such adaptive strategies might enhance the likelihood of reproductive success over the life course, and often involve trade-offs between growth and reproduction, as described and rationalized in life history theory^7–10^. Although this theory explains phenotypic diversity and plasticity in the context of resource availability^11–13^, a mechanistic understanding at the molecular level, detailing how an altered reproductive strategy can be implemented and maintained throughout the life course, is completely lacking.

Epigenetic modification provides a means of sensing the environment and translating diverse signals into altered patterns of gene expression, which can have a profound and long-term effect on the phenotype. The epigenome undergoes considerable modification during various stages of development and, as is becoming clear, plays a role in the maturation of the reproductive axis at puberty as well as adult reproductive function^14–16^. We therefore hypothesized that the early-life environment programs reproductive strategies through epigenetic-driven mechanisms.

Multiple hurdles exist in studying the epigenetic and mechanistic bases to human reproductive function and plasticity, not least of which is the inaccessibility of hypothalamic-pituitary-gonadal (HPG) tissues in healthy subjects^1^. In this study, we first examined buccal tissues from Bangladeshi women to look for changes in DNA methylation associated with their childhood environment. In order to address the functional relevance of these findings, we then employed a mouse model of early-life challenges comparable to those experienced in Bangladesh. The women’s distinct reproductive phenotype is associated specifically with higher disease load in Bangladesh^3,4^, where individuals are exposed to recurrent immune challenge in a country prone to seasonal floods, outbreaks of disease and relatively poor healthcare^17^. The Bangladeshi women in our studies are, however, well-nourished, rarely perform manual work and are relatively affluent. To match these conditions, we adopted a mouse model of pre-pubertal (equivalent of human age ∼6.5-9 y) mild colitis to expose the mice to immunological challenges. The treatment leads to a similarly altered reproductive phenotype and permits access to the functional reproductive tissues for gene expression and epigenetic analysis. This approach, combining all the advantages of an experimental mouse model with observations and proxy tissue DNA analysis from distinct groups of women, has revealed a pivotal role for the steroidogenic enzyme, 5α reductase-1 and its epigenetic regulation in programming adult reproductive strategies, affecting the timing of pubertal onset, adult reproductive function and duration of the reproductive lifespan.

## Results

### Differential methylation patterns in Bangladeshi women who grew up in Bangladesh or in the UK

Methylation analysis of buccal DNA revealed that adult Bangladeshi women (aged 28.1 ± 5.0 y) living in London had distinct methylation signatures, depending on whether they had experienced childhood in Bangladesh (n=15) or the UK (n=13). Illumina-Methylation Epic array data revealed 17,004 CpG sites with a mean methylation difference >20%, most of which (14,509) mapped to “open sea” regions of the genome; a smaller number (2,423) were associated with “shores” and “shelves” and 72 mapped within CpG islands. Methylation of CpG islands harboring promoters is a strong indicator of suppressed gene expression, and we first investigated genes with known functions in fertility. We identified, and confirmed by targeted bisulfite sequencing (Fig S1A), elevated methylation levels in CpG islands associated with *FZD1* and *RUNX3*, both of which encode proteins that regulate ovarian folliculogenesis^18,19^, and also *RASAL3*, which controls a magnitude of inflammatory responses^20^ and has been linked specifically to inflammatory bowel disease^21^. Pathway analysis revealed that genes with differentially methylated CpGs were significantly enriched in the Hippo (FDR 6.99E-12) and PI3K-Akt (FDR 2.83E-07) signaling pathways (Fig S1B-D), both of which play central roles in regulating follicle recruitment and growth^22,23^.

### The altered reproductive phenotype of a mouse model of early life colitis is similar to that of the women who experienced childhood in Bangladesh

In order to determine whether these epigenetic modifications occur also in the reproductive tissues and play a functional role in the altered phenotype, we set up an appropriate animal model. We induced temporal colitis in newly-weaned female mice (22-23 d old: approximately equivalent to 6-6.5 y in human age^24^), by administration of dextran sodium sulfate (DSS) in the drinking water for 7 days, in order to mimic early-life immune challenges experienced by girls in Bangladesh^3,4^. The mice stopped gaining weight during the latter part of the treatment and blood was evident in the feces, but they quickly recovered (Fig 1A). Notably, however, the DSS-treated mice had delayed onset of puberty by an average of 6 days (Fig 1B), corresponding to just over 1.5 y in human lifespan^24^. This was evident only in the treated mice and, as in the women, was not inherited transgenerationally^3^, with puberty in the female off-spring occurring at a similar age to that in off-spring of littermate controls (Fig 1C). Levels of circulating anti-Müllerian hormone (AMH) comprise a clinical marker for size of the ovarian reserve and, as in women who had experienced Bangladeshi childhood^5^, were significantly lower in the DSS-treated mice than the controls (Fig 1D). Thus, the reproductive phenotype of the mouse model mirrors that of the women who were exposed to the early-life immune challenges of childhood in Bangladesh, enabling us to study the underlying mechanisms of this adaptive response.

**Fig 1:**
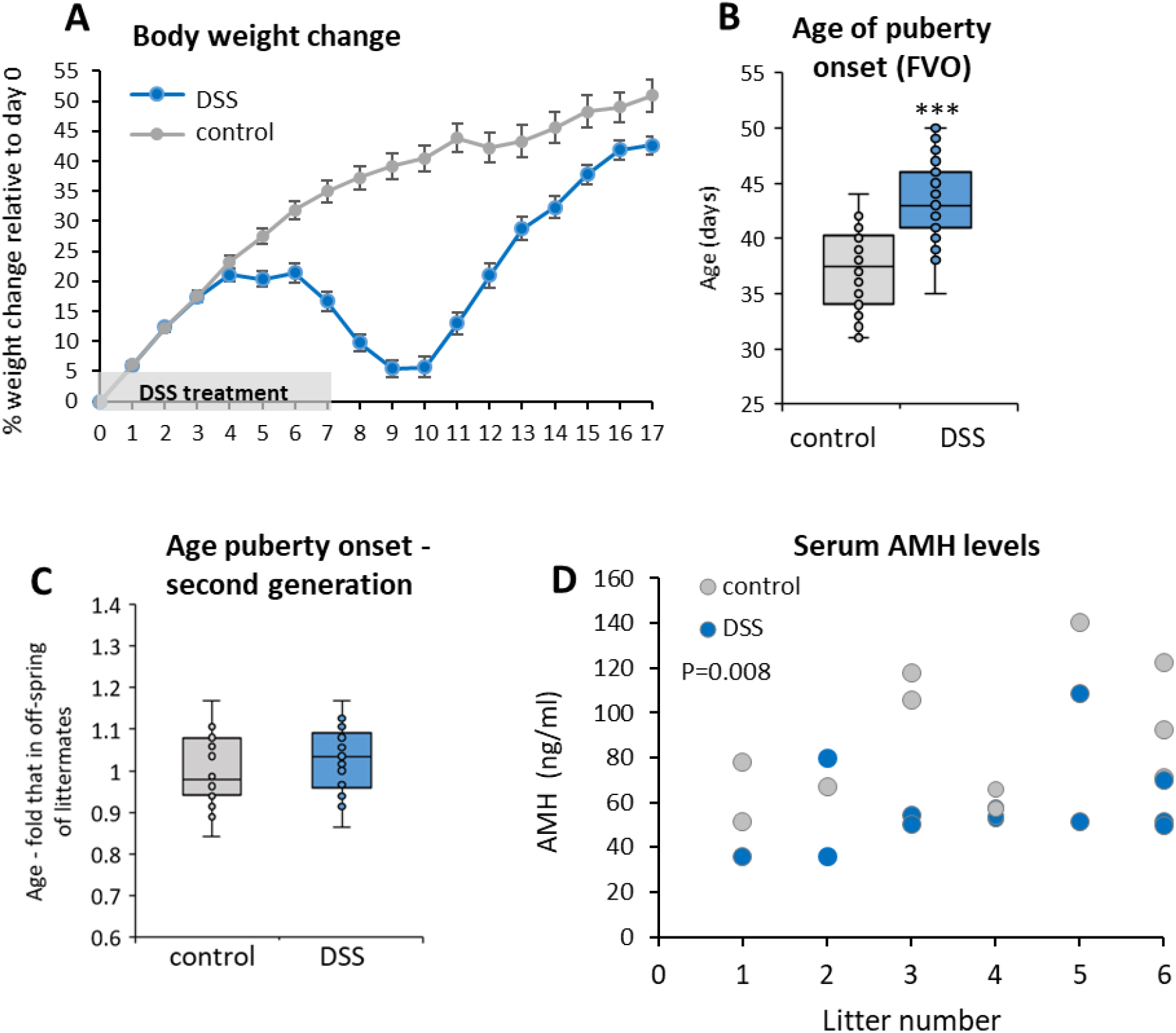
Reproductive phenotype of the mouse model of early-life colitis is similar to that of the women who experienced childhood immunological challenge. (A) Changes in body weight in control (n=48) and DSS-treated (n=45) mice, relative to their initial weight. Mean±SEM. (B) Age of first vaginal opening (FVO), indicating onset of puberty. ***P<0.001; n=44, 48. (C) Age of FVO in second generation, relative to that in off-spring of parent littermate controls. P>0.05; n=20, 30. (D) Circulating AMH (ng/ml) in control and DSS-treated mice, shown separately for each of six litters. Mean between groups (control: 88.07 ng/ml [n=11] and DSS: 57.06 ng/ml [n=14]: P=0.008).

### The mouse model ovaries have altered follicle numbers and gene expression in pathways regulating follicle growth and recruitment

Given that these findings pointed to altered ovarian function, we examined histological sections of the mouse ovaries. Follicle counts showed fewer primary and antral follicles in the DSS-treated mice than in the controls, and there were significantly more atretic follicles (Fig 2A-E). This concurs with the lower AMH levels, and reveals an altered reproductive trajectory involving increased rates of oocyte depletion from the limited ovarian pool.

**Fig 2:**
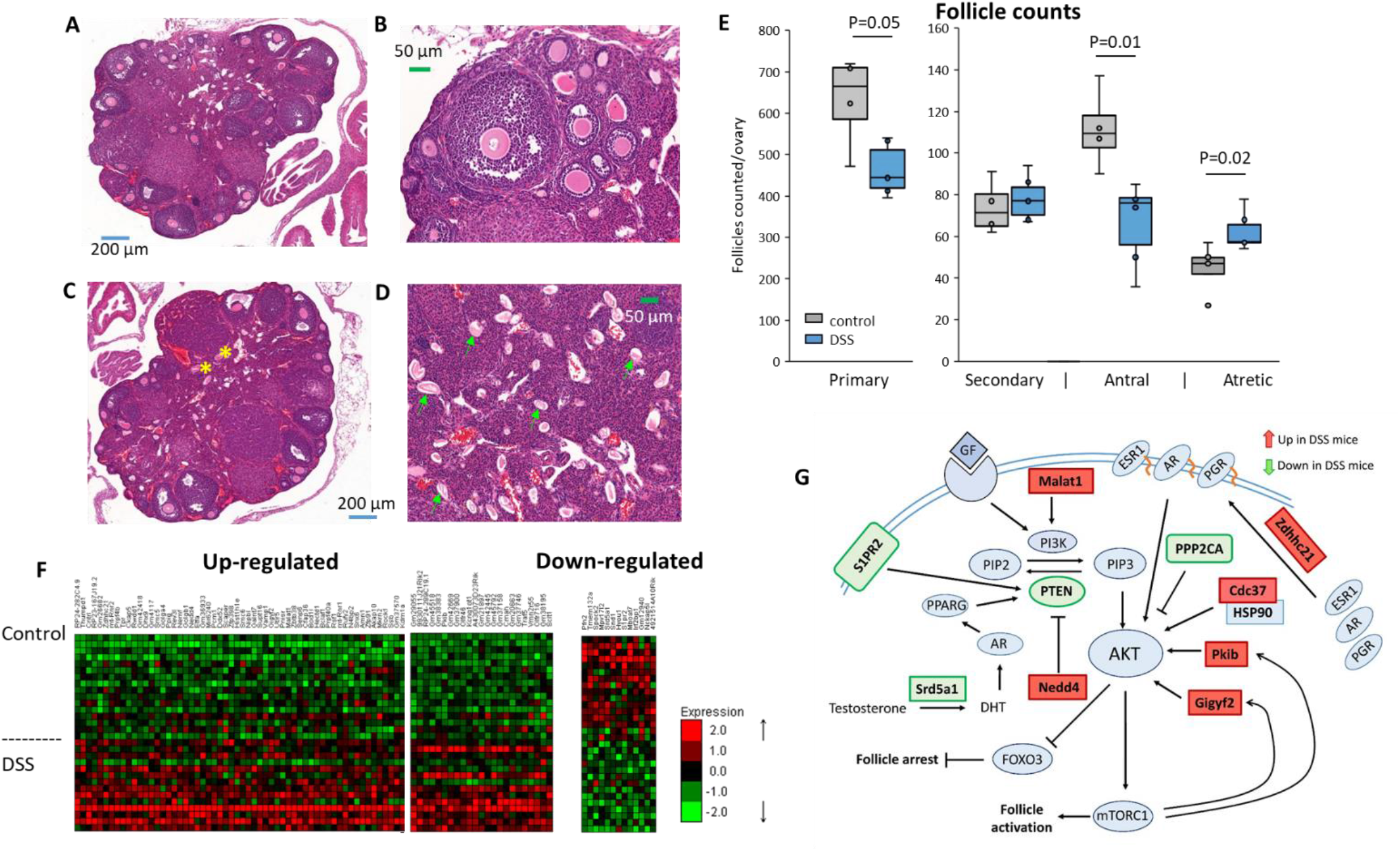
The mouse model ovaries exhibit altered follicle numbers and pathways of gene expression regulating follicle activation. (A-D) H&E stained ovarian histological sections from (A,B) control and (C,D) DSS-treated groups. Some atretic follicles (yellow asterisks) and *zona pellucida* remnants (green arrows) are marked. (E) Follicle counts from sections of mice ovaries (n=4, n= 6), compared by t-test. (F) Heat map of differentially expressed ovarian genes (DEGs) at P*adj*<0.05 from RNA-seq analysis. Each column represents a gene and each row represents the expression level in one ovary. (G) Signaling pathway to follicle activation, showing some of the DEGs (P<0.05). Red boxes signify up-regulated genes; green boxes signify down-regulated genes.

In order to determine the pathways responsible for the altered ovarian activity, we carried out RNA-seq transcriptome analysis of ovaries from the treated mice and their litter-mate controls. Both coding and non-coding RNAs were found to be differentially expressed: 92 were upregulated, while 13 were down-regulated (P*adj* <0.05; Fig 2F). Pathway analysis (Fig S2) of the differentially-expressed genes (P<0.05) revealed enrichment specifically for oocyte-meiosis (FDR 1.9-E02) and, as in the women’s differentially methylated DNA, also for the Hippo signaling pathway (FDR 4.3-E01). Also similar to the women’s distinct methylation patterns, genes in the PI3K-AKT signaling pathway that stimulates the recruitment of ovarian follicles^23^ were enriched, with up-regulation of several activators of the pathway, while the expression of PTEN which represses this pathway was reduced (Fig 2G).

### SRD5A1 is hypermethylated in women and mice following early-life immune challenge

Among the most significantly differentially expressed genes in the mouse ovaries, three were associated specifically with differentially methylated regions in the women’s DNA. *PKIB* and *GIGYF2* (both encode proteins that activate AKT; Fig 2G) were less methylated in the women who experienced childhood in Bangladesh and their expression was up-regulated in the ovaries from the DSS-treated mice, while *SRD5A1* was more methylated in these women and its expression down-regulated in the mouse model (Fig 3A-C).

**Fig 3:**
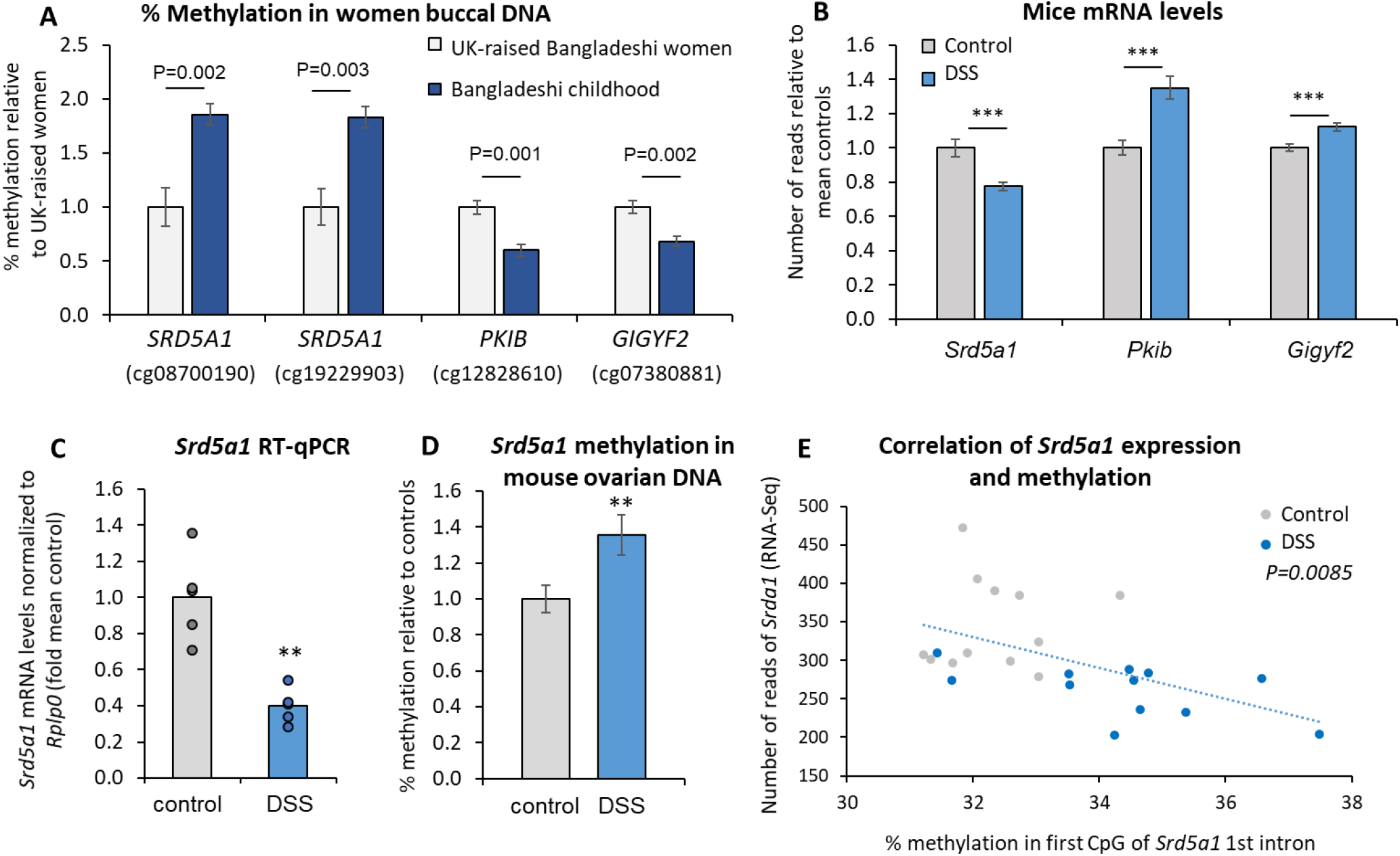
Srd5a1 is hypermethylated in women and mice following early-life immune challenge. (A) Three genes (Illumina EpicMethylation sites) associated with differentially methylated regions in buccal DNA of Bangladeshi women who grew up in Bangladesh (n=16) or UK (n=13); mean±SEM. (B) The mRNA levels of these genes in control (n=16) and DSS-treated (n=14) mice ovaries from the RNA-seq analysis; ***: P*adj*<0.001; mean±SEM. (C) qPCR analysis of the *Srd5a1* mRNA levels (n=5), **: P=0.007; showing means with individual data points. (D) Levels of CpG methylation in the 5’region of the *Srd5a1* first intron (corresponds with first site in Fig 3A: see S3), in control (n=24) and DSS-treated (n=29) mice ovaries, mean±SEM shown relative to controls; **: P=0.015 (Mann-Whitney t-test). (E) Correlation between the levels of *Srd5a1* mRNA (from RNA-seq analysis) and methylation measured in the same samples; P=0.0085.

*SRD5A1* encodes the steroidogenic enzyme, 5α reductase-1, which converts testosterone to dihydrotestosterone (DHT). DHT inhibits follicle activation through decreasing cyclin D2 expression and inducing cell cycle arrest^25^, and via activation of PTEN which represses PI3/AKT signaling^26^. DHT also activates progesterone production^27^. Thus the drop in 5α reductase1 levels would not only facilitate oocyte exit from the primordial follicle pool, in accordance with the ovarian histology, but would also lower progesterone production, as seen in the women who spent their childhood in Bangladesh^3^.

We therefore examined whether methylation of *Srd5a1* was also affected in our mouse model. The *Srd5a1* promoter was reported to be hypermethylated in the prefrontal cortex of mice in response to psychological stress^28^, but it was completely unmethylated in ovaries of both DSS and control groups of mice (Fig S3A). Only ∼570 bp separate the start of the mouse *Srd5a1* gene and its neighboring gene, *Nsun2*, while in the human genome, these divergent genes are just ∼50 bp apart, which points to additional key gene-specific *cis* regulatory regions beyond the proximal promoter. We therefore examined the mouse genomic region, homologous to that differentially methylated in the human samples, which is located in both genomes at the start of the first intron. DNA bisulfite conversion and deep-sequencing revealed that this region was significantly more methylated in the ovaries of the DSS-treated group than in the controls (Fig 3D). This intronic region carries all of the marks of a transcriptional enhancer^29^ (Fig S3B), supporting the impact of this methylation on gene expression levels, which was observed also in the strong correlation between levels of mRNA expression and CpG methylation at the locus (Fig 3E). Furthermore, this putative enhancer region contains a SNP, rs494958, which was reported to be associated with age at natural menopause, as well as two other significant SNPs in high linkage disequilibrium with this trait^30^.

### The up-regulation of Srd5a1 by estradiol is blunted by anti-inflammatory cytokines

To determine the pathways leading to a reduction in *Srd5a1* expression as a result of the early-life adversity, we first examined how its levels normally change with sexual maturation. Comparison of *Srd5a1* mRNA levels in ovaries from sexually immature and mature mice showed that they increased around 6-fold over this time (Fig 4A). To determine the mechanisms involved, we exposed ovarian KK-1 mouse granulosa cells to various steroids, which revealed a stimulatory effect of estradiol (E2), while neither DHT, progesterone nor dexamethasone (synthetic cortisol) had any notable effects (Fig 4B, S4). This suggests that the rise in *Srd5a1* expression at the time of puberty is due to the increase in E2 levels.

**Fig 4:**
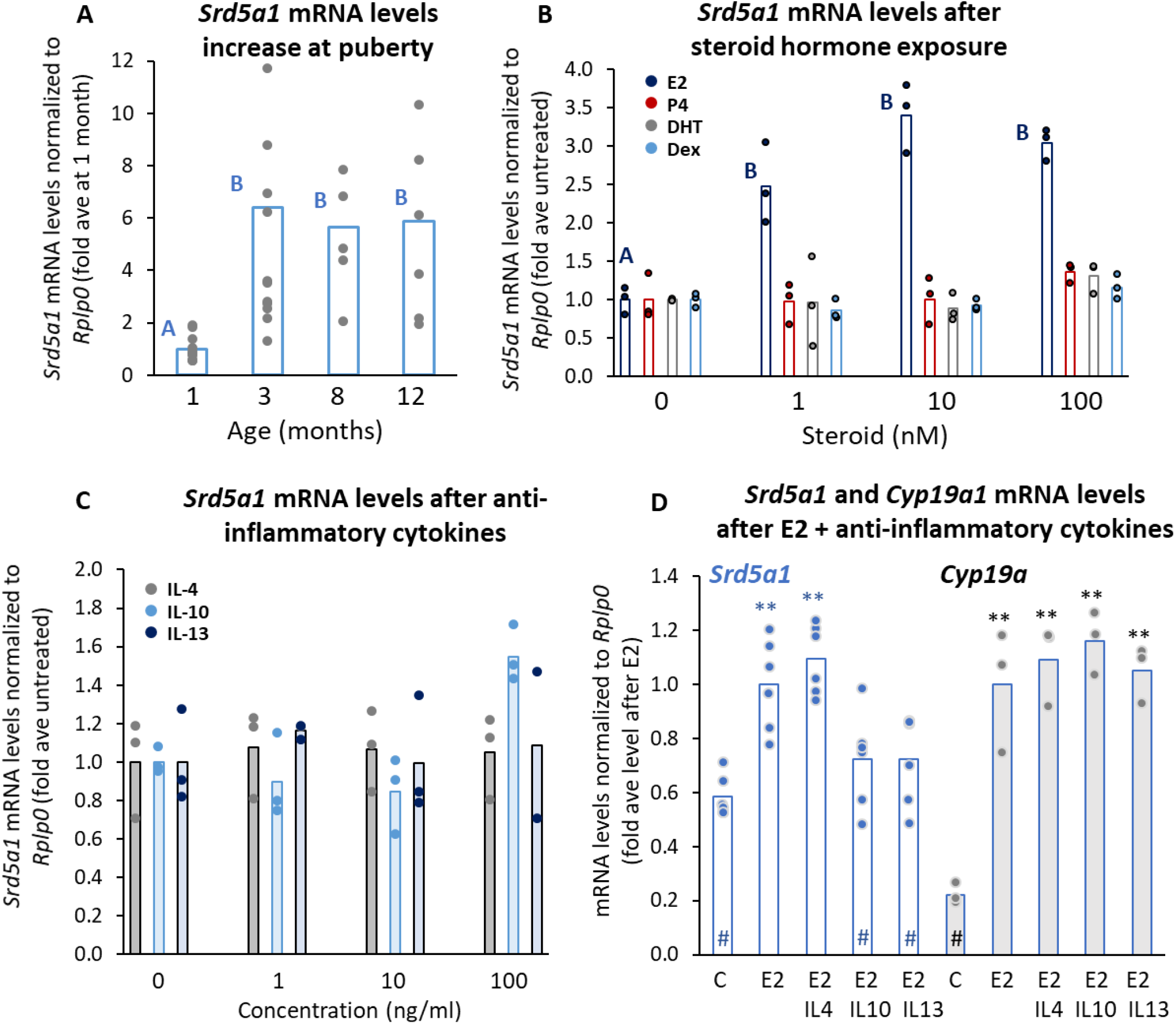
The up-regulation of Srd5a1 by estradiol is blunted by anti-inflammatory cytokines. (A) *Srd5a1* mRNA levels in ovaries of mice of various ages (n=12, 14, 5 or 6); for groups sharing same letter: p>0.05 (ANOVA, Tukey-Kramer t-test). (B) *Srd5a1* mRNA levels in KK-1 cells (n=3) after exposure to estradiol (E2), progesterone (P4), dihydrotestosterone (DHT) or dexamethasone (Dex). For E2, ANOVA is as in Fig 4A; otherwise P>0.05 (C) *Srd5a1* mRNA levels after cytokine exposure (n=3). (D) *Srd5a1* (n=6) and *Cyp19a* (n=3) mRNA levels after E2 alone (10 nM) or with cytokine (100 ng/ml); **: p<0.02 vs control; #: p<0.02 vs E2; where not marked p>0.05. All graphs show mean with individual data points.

We then verified whether *Srd5a1* expression is affected adversely by the anti-inflammatory cytokines, IL4, IL-10 and IL-13, which are elevated in the general stress response. Although these cytokines alone did not reduce basal *Srd5a1* levels (Fig 4C), when given together with E2, both IL-10 and IL-13 blocked the E2-stimulatory effect on this gene without affecting its up-regulation of *Cyp19a1* (Fig 4D). Thus, early life adversity leading to an increase in IL-10 and/or IL-13, can cause a reduction in 5α reductase-1 levels by dampening the stimulatory effect of E2, explaining the particularly significant impact of this response at early stages of pubertal development.

### 5α reductase-1 regulates the central control of reproduction and pubertal timing

5α reductase-1 is widely expressed and we considered that its altered expression in non-ovarian tissues might play additional roles in mediating the distinct reproductive phenotype. We therefore measured *Srd5a1* expression in the hypothalamus and the prefrontal cortex of the brain, and in the pituitary of the mouse model. *Srd5a1* mRNA levels were reduced in the hypothalamus of the treated mice, but not in the prefrontal cortex or the pituitary (Fig 5A). Using additional mice, separation of the hypothalamus into distinct regions confirmed the reduced expression of *Srd5a1* specifically in the pre-optic area (Fig 5B), which contains most of the neurons that control reproduction, suggesting a possible role in the timing of pubertal onset. This connection was supported by the fact that in second generation mice, both age of pubertal onset (Fig 1C) and levels of *Srd5a1* mRNA in the ovaries and hypothalami (Fig 5C) were similar to those of controls.

**Fig 5:**
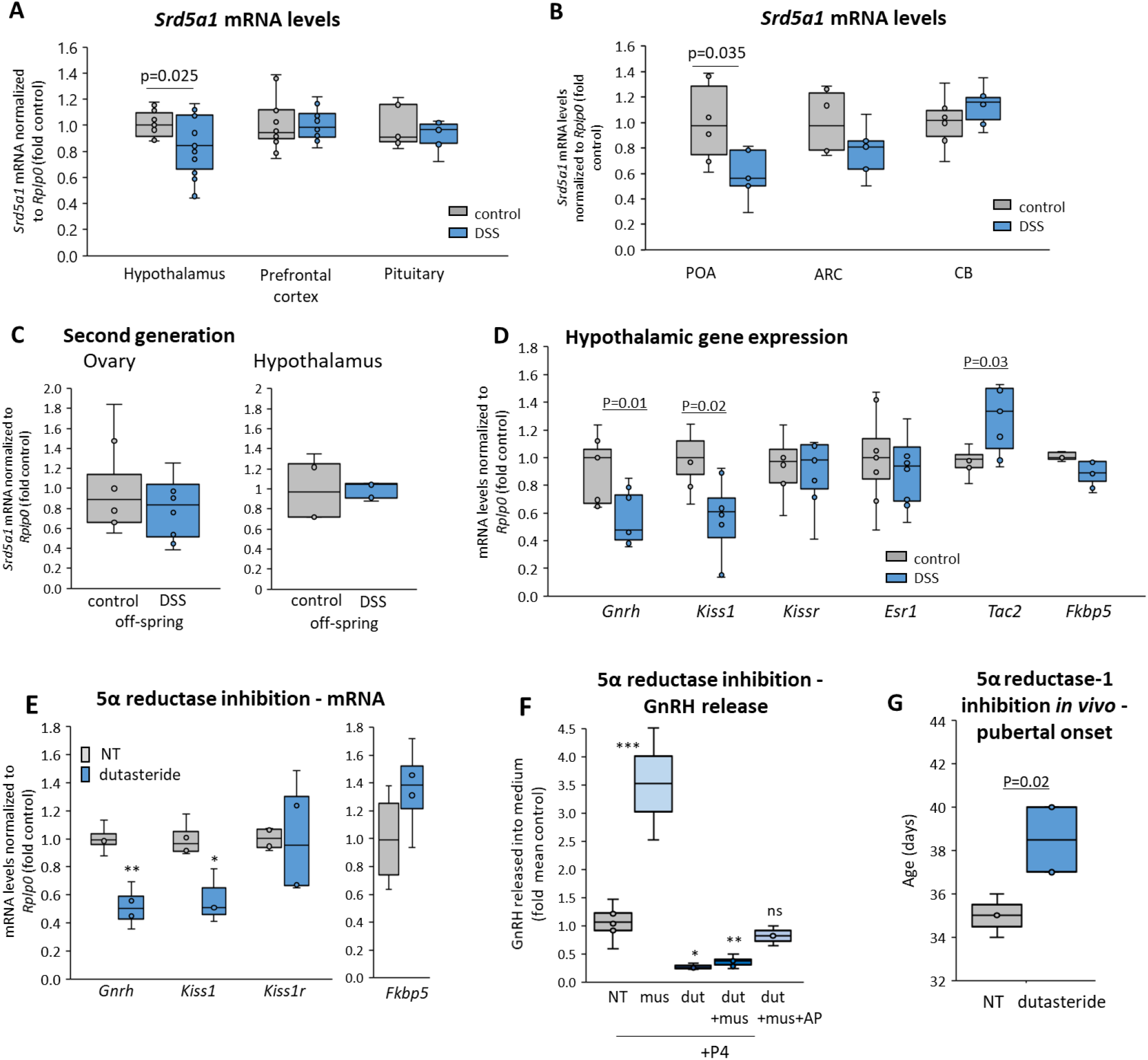
5α reductase-1 regulates the central control of reproduction and pubertal timing. *Srd5a1* mRNA levels were measured in (A) hypothalamus (n=14), prefrontal cortex (n=15,14) and pituitary (n=7) of control and DSS-treated mice; or (B) preoptic area (POA: n=6, 5) and arcuate nucleus (ARC: n=6, 5) of the hypothalamus, and the cerebellum (CB: n= 6). (C) *Srd5a1* mRNA levels in the ovary (n=8) and hypothalamus (n=4,5) of female off-spring of DSS-treated mice and their littermate controls. (D) The mRNA levels of genes encoding reproductive regulatory factors were measured in the hypothalamus of the DSS-treated and control mice, with *Fkbp5* as an indicator for stress. (E,F) The effect of 5α reductase inhibitor, dutasteride in GT1-7 GnRH neuronal cells on (E) gene expression (n=8, 4) and (F) GnRH release, in which some cells were also exposed to the GABAA agonist, muscimol, alone (n=2), with dutasteride (n=6) or together with AP (n=3), all in the presence of AP precursor, P4. Significant differences (p<0.05) are shown for comparisons with control mice or untreated cells, otherwise P>0.05. (G) Age of FVO following dutasteride: female mice and their control littermates were given dutasteride (or vehicle) in the diet each day after weaning, and checked daily for FVO (n=3, 4).

The drop in *Srd5a1* expression in the hypothalamus led us to examine the expression levels of *Gnrh* and other genes that encode factors regulating the reproductive axis. The mRNA levels of *Gnrh* were significantly lower in the DSS-treated mice than in their litter-mate controls, as were those of *Kiss1*; notably *Tac2* mRNA levels were elevated, while those of *Kiss1r* and *Esr1* were unaltered (Fig 5D). The expression of *Fkbp5*, which is highly sensitive to glucocorticoids, was not different in these mice (Fig 5D), indicating that the mice were not suffering chronic stress.

To establish whether 5α reductase-1 activity regulates expression of these genes in GnRH neurons, we exposed the GT1-7 GnRH neuronal cell line to the inhibitor, dutasteride. As in the DSS-treated mice, both *Gnrh* and *Kiss1* mRNA levels were repressed by the drop in 5α reductase-1 activity, while the mRNA levels of *Kiss1r* and *Fkbp5* were unaffected (Fig 5E). Given previous reports that some neurosteroids catalysed by this enzyme activate the stimulatory GABA-A receptor and can augment GABA effects on GnRH release at puberty^31–33^, we also examined the effects of 5α reductase-1 inhibition on GnRH release. Dutasteride repressed GnRH release from these cells and completely blocked the stimulatory effects of a GABA-A agonist, muscimol; however, levels were restored to those of controls by addition of the neurosteroid, allopregnanolone (Fig 7F).

Having established the impact of early-life adversity on hypothalamic expression of *Srd5a1*, and the effects of 5α reductase-1 on regulation of GnRH synthesis and secretion, we went on to demonstrate the role of this enzyme in determining pubertal onset *in vivo*. Young female mice were treated with dutasteride daily starting soon after weaning. The treatment delayed their first vaginal opening (FVO) by 3-4 days compared to the sham-treated litter-mate controls (Fig 7G), which corresponds to an estimated ∼1 y in human lifetime^24^. Thus the reduced expression of this enzyme due to down-regulation of *Srd5a1* expression following early-life adversity appears to play a role in the delay in pubertal onset.

## Discussion

Our study demonstrates that down-regulation of *SRD5A1* plays a pivotal role in shaping adult reproductive function in response to experiences during pre-pubertal development. Having found that this gene was differentially-methylated in buccal DNA of Bangladeshi women according to their childhood environment, we have been able to confirm, in an appropriate mouse model, its altered expression in the functional reproductive tissues and its function in the distinct reproductive phenotype that follows this early-life adversity. Through manipulations *in vitro* and *in vivo*, we have demonstrated the role of 5α reductase-1 in determining pubertal timing through regulation of GnRH synthesis and secretion. Moreover, the modified expression in the mouse model of *Srd5a1* and other genes in the pathways to ovarian follicle recruitment, explains the differing rates of oocyte depletion and disparate ages at menopause in the Bangladeshi women. We have thus uncovered a key role for this enzyme in determining the reproductive trajectory throughout the adult lifespan, from puberty to menopause, which is established by epigenetic programming during childhood development.

Given that *Srd5a1* mRNA levels increase across puberty and in response to E2, the immediate pre-pubertal period evidently comprises a particularly sensitive time to any signals that diminish this up-regulation. Such signals include the anti-inflammatory cytokines that we examined, and likely encompass additional signals arising from other forms of physiological or psychological stress, given that activity of this enzyme was seen to be decreased following various stressors and this was most pronounced when the stress was experienced at a young age (e.g.^34–38^). These reports support a role for epigenetic modification of *SRD5A1* during childhood in determining additional aspects of the adult phenotype, particularly relating to the stress axis.

Apart from its role in pubertal timing, reduced expression of 5α reductase-1 in the hypothalamus undoubtedly has consequences for other endocrine axes, especially those controlling growth and the stress response which are also regulated by neurosteroids catalysed by this enzyme^38–40^. In the face of adversity and limited resources, altered epigenetic regulation of *SRD5A1* in the hypothalamic control center, would allow differential regulation of these major endocrine axes to mediate the adaptive response to changing environmental conditions which underlie the trade-offs between growth, reproduction and homeostasis described through life history theory.

Such reprogramming of these axes might well be beneficial for the individual, but the altered reproductive phenotype presents health consequences, given that timing and duration of the reproductive lifespan dictate risks for steroid- and age-related disease. The epigenetic basis of plasticity that we describe here explains some of the diversity in reproductive characteristics and how they are shaped by early childhood environment, while also opening up the possibility that these characteristics might be susceptible to manipulation to mitigate health issues across the lifespan.

## Methods

### Human methylation analysis

British-Bangladeshi women (20-35 y), were recruited in London through community contacts using snowballing techniques. The first group comprised women who were born in Bangladesh and moved to the UK when aged over 16 y. The second group comprised women who were second-generation British-Bangladeshis, born in the UK to Bangladeshi migrant parents. Protocols for human data collection were approved by the Ethical Committee of the Department of Anthropology, Durham University. Women gave informed consent to participate in the study, and the data were anonymized at source.

Buccal swabs were collected with iSWAB (Mawi DNA Technologies) and genomic DNA isolated using the DNeasy Blood & Tissue Kit (Qiagen). Genome-wide DNA methylation data acquisition was carried out on the Infinium MethylationEPIC BeadChip platform (Illumina) and performed by Tepnel Pharma Services, UK using ‘Bangladeshi childhood’ (n=16) and ‘UK childhood’ (n=13) DNA samples, which passed the quality control checks. Multidimensional scaling (MDS) plots indicated that no significant batch effects were skewing our MethylationEPIC BeadChip data sets. The data were processed with the Bioconductor/minfi package. CpG probes associated with known SNPs were removed, as were those with a detection probability of <0.01. Probes on both X and Y chromosomes were retained. Methylation beta values (0-1) were normalized by SWAN. Dmpfinder/minfi was applied to determine probes with significantly differentially methylation levels between the ‘Bangladeshi-childhood’ and the ‘UK-childhood’ groups. FDR was set at <0.05. The pathway analysis utilized NIPA, a tool that performs enrichment tests by hypergeometric statistics (https://github.com/ADAC-UoN/NIPA/).

For validation of the array data, targeted bisulfite sequencing was conducted on a subset of the samples used to generate the MethylationEPIC BeadChip data (chosen on the basis of their DNA quality and concentration) via amplicon sequencing of 10 CpG sites using the MiSeq system (Barts and the London School of Medicine and Dentistry, GenomeCentre). The interrogated CpGs were cg25470148 (chr1:25257931) lower-strand,*RUNX3*; cg16696646 (chr17:19861616), lower-strand *AKAP10*; cg26916966 (chr17:40274524), upper-strand, *KAT2A*; cg07357279 (chr17:43318735), upper-strand, *FMNL1*; cg01062942 (chr19:15568935), upper-strand, *RASAL3*; cg08470875 (chr2:26401718), upper-strand, *FAM59B*; cg08700190 (chr5:6636046), upper-strand, *SRD5A1*; cg08198075 (chr6:123033536), upper-strand, *PKIB*; cg12914114 (chr6:170687002), lower-strand, *FAM120B*; cg01480180 (chr7:90896329), lower-strand, *FZD1*. For each interrogated genomic region, >100 sequencing reads were obtained. Primers are given in Table S1.

### Mice

All animals were held and handled humanely, after protocol approval, and in accordance with IACUC guidelines. For DSS-treatments, upon weaning, female mice from each litter were divided randomly into two groups to provide littermate controls for all experiments. After ∼2 d recovery, the mice were ear-marked and weighed, and one group received 3 % dextran sodium sulfate (DSS: 35-50 kDa, MP Biochemicals) in the drinking water for 7 d. The DSS water was changed every 2-3 d and the mice were weighed each day for at least 16 d. All mice were observed daily for signs of pubertal onset, as determined by FVO. In order to assess the impact of this early life exposure on the second generation, a single male was housed with the DSS-treated and the littermate control female mice.

For harvest of brain tissue, the brains were removed and whole hypothalamus or prefrontal cortex isolated into 1 ml TRIzol for RNA extraction. For isolation of specific regions (preoptic area, arcuate nucleus or cerebellum), brains were transferred into a brain matrix (RWD-800-00149-00) for coronal sectioning following isolation of each region from the relevant section, as determined using Allen Brain Atlas. All tissues were collected from females in estrous, verified by cytological smears. Blood was collected by cardiac puncture at the time of sacrifice, and circulating AMH levels were measured by ELISA (Ansh labs, Webster, Texas) according to the manufacturer’s protocol, after dilution of all samples x20.

For the administration of dutasteride *in vivo*, after weaning, female mice from each litter were marked, weighed and divided to two groups. A transparent plastic separator (kindly given by Madaf Plazit Packaging) was inserted into each cage, with one mouse in each half. Dutasteride (SML1221, Sigma) was dissolved in oleic acid (O1383, Sigma) at 15 mg/ml, and added to a ∼60 mg piece of enriched diet pellet (D12451i, Research Diets). Immediately after separation, each mouse received the dutasteride-treated diet (∼13 μg/g BW) or a similar amount of vehicle-treated diet for controls. The pellet was consumed fully within a few minutes, and the separator was then removed. The treatment was repeated daily, and mice were weighed and observed daily for signs of FVO.

### Histology and follicle counts

Ovaries were harvested from the mice at ∼60 d and were fixed with 4 % paraformaldehyde for 4 h before transferral to 70 % ethanol. Paraffin embedding, sectioning (4 μm), and hematoxylin and eosin (H&E) staining were carried out at the Biomedical Core Facilities at the Rappaport Faculty of Medicine, Technion-Israel Institute of Technology. Identification of follicle stage (using CaseViewer software) and counting were performed (as in^41,42^), while blind to the treatment group. In short, every fifth section per ovary was analyzed, and the follicular stage was determined by size and morphological characteristics: primary follicles containing a single layer of cuboidal granulosa cells; secondary follicles showing more than one layer of granulosa cells but no antrum, and antral follicles containing an antral space. Atretic follicles were identified based on the presence of *zona pellucida* remnants, stained bright pink. Secondary and antral follicles were counted only if a nucleus was present, and the atretic follicles, which vary considerably in size, were counted every 8^th^ stained section, to avoid counting the same follicle twice. The “follicle counts” presented comprise the number of follicles at each of these stages counted in each ovary using this approach.

### Quantitative PCR, transcriptome and methylation analysis

RNA was isolated using TRIzol, DNase I-digested and cleaned using R1014 RNA Clean & Concentrator-5 kit (Zymo Research), cDNA synthesized using the qScript Flex cDNA kit (95049 Quanta) using oligo dT, and real-time quantitative PCR (qPCR) was carried out using the PerfeCTa SYBR Green FastMix (Quanta), both as previously reported^43^, using primers listed in Table S2. Amplicon levels were quantified using standard curves and normalized to levels of *Rplp0*.

For transcriptome analysis, RNA was extracted from the mice ovaries and purified as above, and sequenced by CEL-seq, using Illumina HiSeq 2500 at the Technion Genome Center (as in^44^). FastQC was used for quality control and the reads were mapped by TopHat algorithm to mm10 genome assembly. HTSeq-count was used to count the reads, and the normalization of raw counts and differential expression were calculated using DESeq2 in R platform, with P*adj* using Benjamini and Hochberg correction for false discovery. Pathway analysis was performed using the Database for Annotation, Visualization and Integrated Discovery (DAVID).

DNA was extracted from the mouse tissues using TRIZOL, and the genomic DNA cleaned with the Quick-DNA Miniprep Plus Kit (D4068; Zymo), before bisulfite conversion using the EZ-DNA Methylation-Gold Kit (D5005 Zymo), and two rounds of PCR-amplification (nested, with outer and inner primers: Table S2) using Red Load Taq Master (Larova). After purification of the amplicons (PCR purification kit; Qiagen) and cloning into pGEM-T-easy, inserts from 7-8 randomly selected clones were sequenced and analyzed as previously^45^. Subsequently, the region in the first intron homologous to that differentially methylated in the human samples, was amplified and cleaned as above. Additional rounds of PCR were then performed using KAPA HiFi HotStart Ready mix (Roche), initially with primers containing the adaptors (Universal adaptors; Illumina) and subsequently another 8-12 PCR cycles with the specific primers (Illumina Nextera XT index kit); samples were cleaned with PCR purification kit (Qiagen) between each stage. These libraries, after addition of 50% Phi-X, were then deep-sequenced by 150 bp paired-end sequencing on Mi-seq (Illumina), at the Technion Genome Center.

### Cell culture

The KK-1 granulosa cell line was cultured as reported^46^, maintained at 37 °C with 5 % CO_2_ at 30-80 % confluency, passaging 2-3 times a week. The cells were exposed either to 1-100 nM of the steroids (Sigma) for 24 h, or to anti-inflammatory cytokines: IL-4, IL-10 or IL-13 (1-100 ng/ml for 24h), alone or before addition of E2. Alternatively, the GT1-7 mouse hypothalamic GnRH neuronal cell line was cultured with high glucose DMEM containing 10 % FBS, 1 % penicillin-streptomycin, sodium pyruvate and sodium bicarbonate, maintained at 37 °C with 5 % CO_2_ at 50-90 % confluency, passaging 1-2 times a week. For mRNA measurements, the cells were cultured in charcoal-stripped FBS medium for 24 h before some were exposed to dutasteride (5 μM) for 24 h. Cells were then harvested for RNA extraction and qPCR analysis as before.

For analysis of GnRH release, cells (in 6-well plates) were washed twice in medium without FBS, incubated in the same medium, and some exposed to dutasteride (10 μM). After 30 min, P4 (2 μM) was added for 5 h, and then muscimol (100 μM) or/and AP were added for 1 h. The medium was collected into 1.5 ml tubes, centrifuged for 2 min at 3000 g and kept at −80 °C for measurement of GnRH in the supernatant by ELISA (Phoenix Pharmaceuticals, catalog # EK-040-02CE), according to the manufacturer’s protocol.

### Statistical analysis

All data are from multiple biological repeats (n-value) which were assayed individually. Results are shown as box plots (whiskers show minimum and maximum values, boxes represent 25–75% data ranges, horizontal lines within boxes indicate the median values), individual values, or as mean ± SEM values. Statistical analysis for parametric data was using a Student’s *t*-test (two-tailed), and differences considered significant at *p* ≤ 0.05, or alternatively One-way analysis of variance (ANOVA), followed by the Tukey-Kramer or Bonferroni *t*-test for multiple comparisons. Methylation analysis (% methylation) utilized Mann-Whitney non-parametric t-test.

### Data Availability

Data are available at NCBI’s Gene Expression Omnibus (GEO), GSE133355 (human DNA methylation data) and GSE133633 (mouse RNA-seq data).

## Acknowledgments

This research was supported by Biotechnology and Biological Science Research Council (BBSRC)/Economic and Social Research Council (ESRC) grant ES/N000471/1 (to GB, RS and PM). We thank Kamila Derecka for technical support.

