## Supplementary Information for "Early-life environment programs reproductive strategies through epigenetic regulation of *SRD5A1*"

**Table S1: Primers for human DNA analysis**

| **Island / shore** | **Gene_Name** | **Strand** | **forward primer** | | **reverse primer** |
| --- | --- | --- | --- | --- | --- |
| chr5:6632740-6634162 | *SRD5A1* | Upper | AAGGGTTAGAGTTATTTTGAATGATAG | TTTCATCCCAACAACTCCTTATCCAA | |
|  | *AKAP10* | Lower | TGGTGAAATTTTGGTTAGAGGT | CCAAAATCCTCCAATCTCTTATCAA | |
|  | *PKIB* | Upper | ATGATTTGTTTTTGTTGGTATATAA | CCCTTATACTACATATTTTCAATTCTCTAC | |
| chr7:90893567-90896683 | *FZD1* | Lower | TGGTATTAGAGTGTGAGGGTAAGAG | CCTTTAAACAACTCAACTCCTACAA | |
| chr6:170686006-170687003 | *FAM120B* | Lower | ATTGTTTTGTTTTAAGGGTTTTTTTGTTA | CCTCCCTAATCCCAAATACCTAAA | |
| chr2:26401695-26402099 | *FAM59B* | Upper | GGTTTTGGAGGGGTGGTG | ACCTTCCCCCTCCTAAAAACCTCTA | |
| chr17:43318429-43319243 | *FMNL1* | Upper | AAGGAGTTTGTTGGTGGGTATT | TCCAACTCCTCCACCTTCAAC | |
| chr17:40274523-40275360 | *KAT2A HSPB9* | Upper | GTTGGGGGATTAATTTGTTGTT | AACTACTCTCCTCCCAAACTCC | |
| chr19:15568027-15569227 | *RASAL3* | Upper | TGGGGGTTTTAGGGTATATAAGAGTAG | CAAAACTCCCCTTCCATCTATTACCA | |
| chr1:25255527-25259005 | *RUNX3* | Lower | TTTTAGAGTTTAGGGAGGTGTTT | CTTCCTCTCCCCCCTCCTAAATCTAT | |

**Table S2: Primers for mice gene and DNA analysis**

| **Gene ID** | **F sequence (5' -> 3')** | **R sequence (5' -> 3')** |
| --- | --- | --- |
| *Rplp0* | GCGACCTGGAAGTCCAACTA | ATCTGCTTGGAGCCCACAT |
| *Srd5a1* | GAATATGTATCTTCAGCCAAC | GGTAATCTTCAAACTTCTCG |
| *Cyp19a1* | CAGTGGAGAGGAGACACTC | CTTCCACCATTCGAACAAGAC |
| *Rasd1* | CTACCATCGAGGACTTCCAC | GACAGGACTTGGTGTCTAGG |
| *Srd5a1* BS outer | AAGGAGTTTTTAGTTAATGTGTGTAG | AAACACAAACTAACACCACCAAAA |
| *Srd5a1 BS* inner | GGAGGTGTTATGTGAAAAATGTTT | CCAAATATCACAAAACTCAACTTC |
| *Gnrh* | GATCCTCAAACTGATGGCCG | CTCCTCGCAGATCCCTGAG |
| *Kiss1* | AGCTGCTGCTTCTCCTCTGT | AGGCTTGCTCTCTGCATACC |
| *Kiss1r* | GGTCATCTACGTTATCTGCC | GCACATGAAGTCTCCCAGC |
| *Esr1* | CTGTGCCGTGTGCAATGACT | CATGCCCACTTCGTAACACT |
| *Tac2* | GTCTCTCTGGAAGGATTGCTG | GGTGTTCTCTTCAACCACGTC |
| *Fkbp5* | GAGTCCAAAGCCTCAGAGTC | GCCAACACCTTCTCGAAGTC |

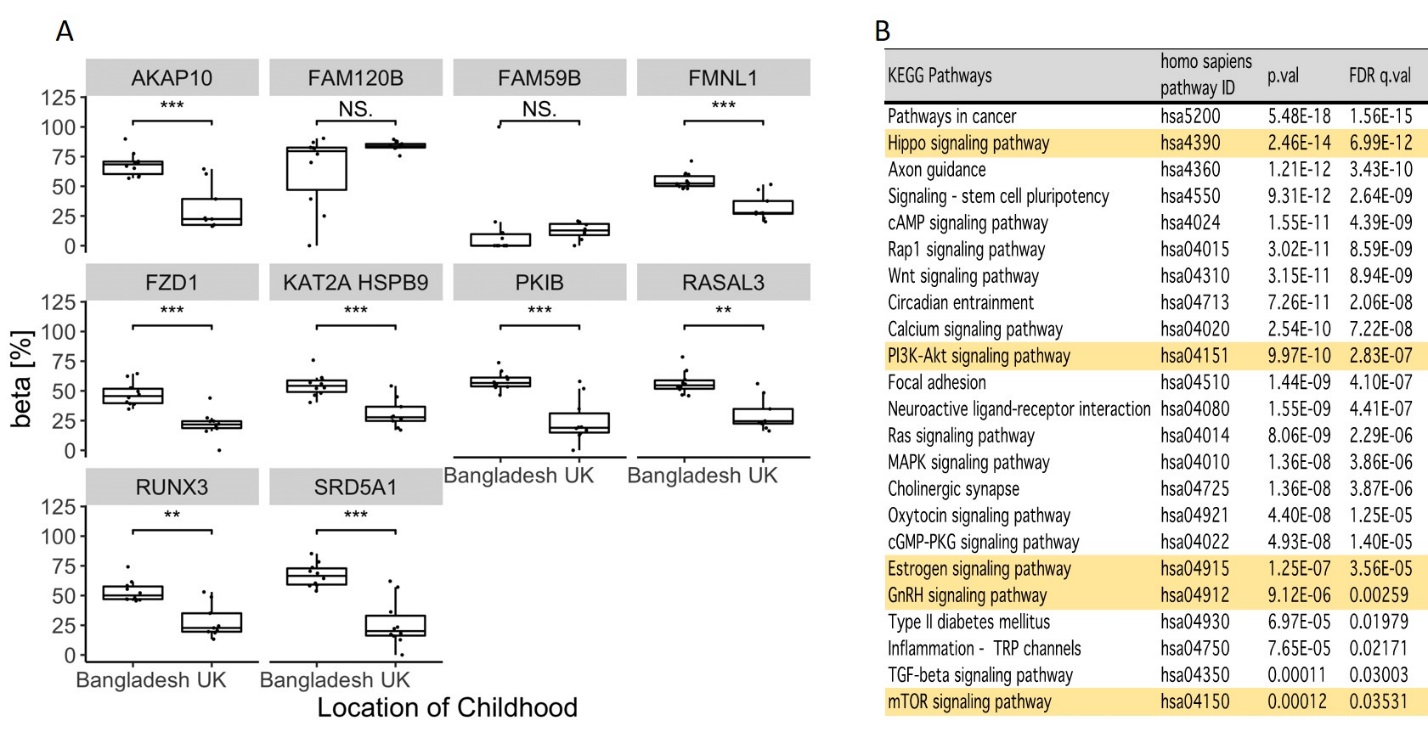

C

D

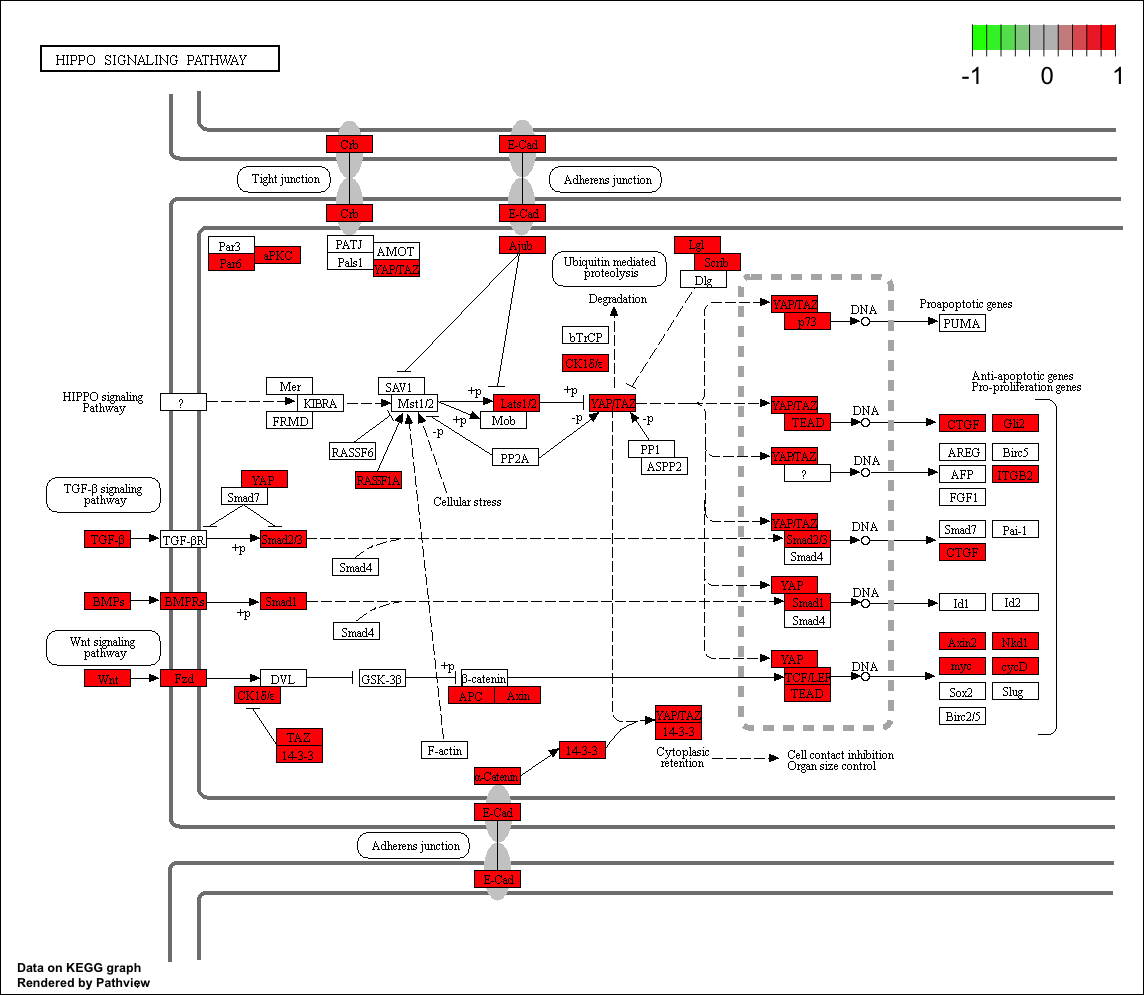

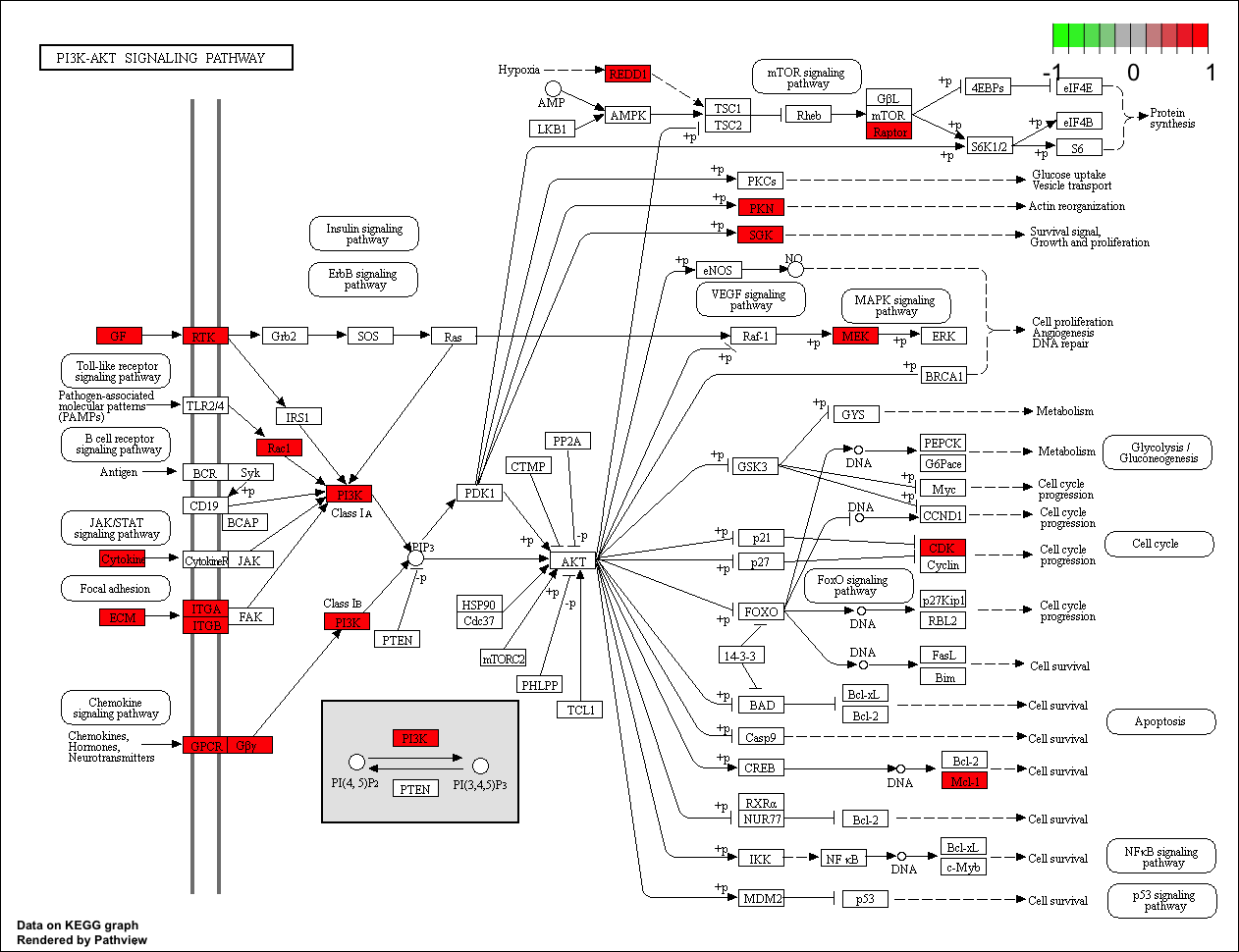

**Fig S1:** *Human Methylation analysis*. (A) Targeted bisulfite sequencing confirmation of MethylationEPIC BeadChip data. Beta values: M/(M+U), where M is number of methylated and U un-methylated reads. Bangladesh-childhood (n=10) and UK-childhood (n=9). Mann-Whitney-U test: *** p<0.001; ** p<0.01; * p<0.05; N.S. p>0.05. (B) KEGG pathway and gene set enrichment analysis of genes associated with differentially methylated probes (FDR <0.05) within islands, shores and shelfs. Pathways of particular relevance for our study are highlighted in yellow. (C,D) Genes in the (C) PI3K-Akt (hsa04151) and (D) Hippo (hsa04390) signalling pathways associated with differentially methylated CpGs in islands, shores or shelfs are highlighted in red.

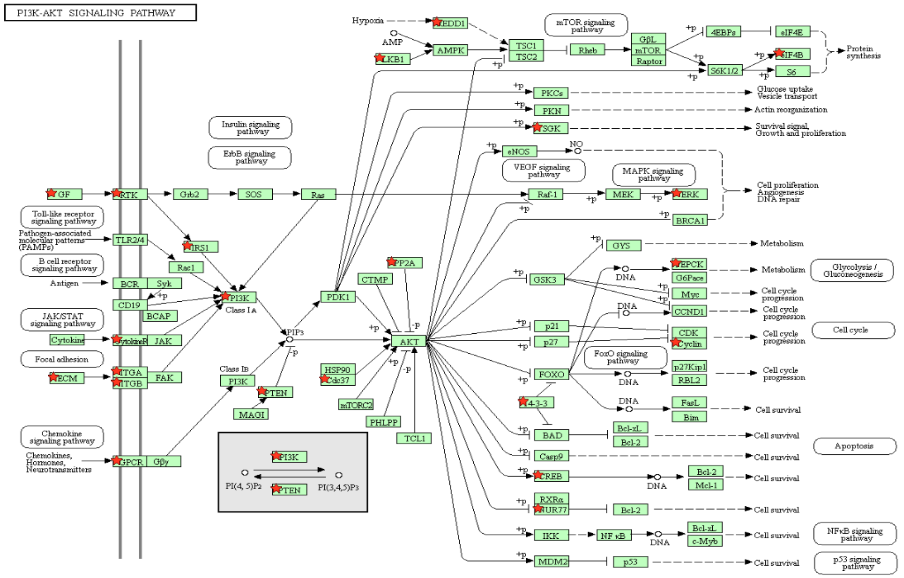

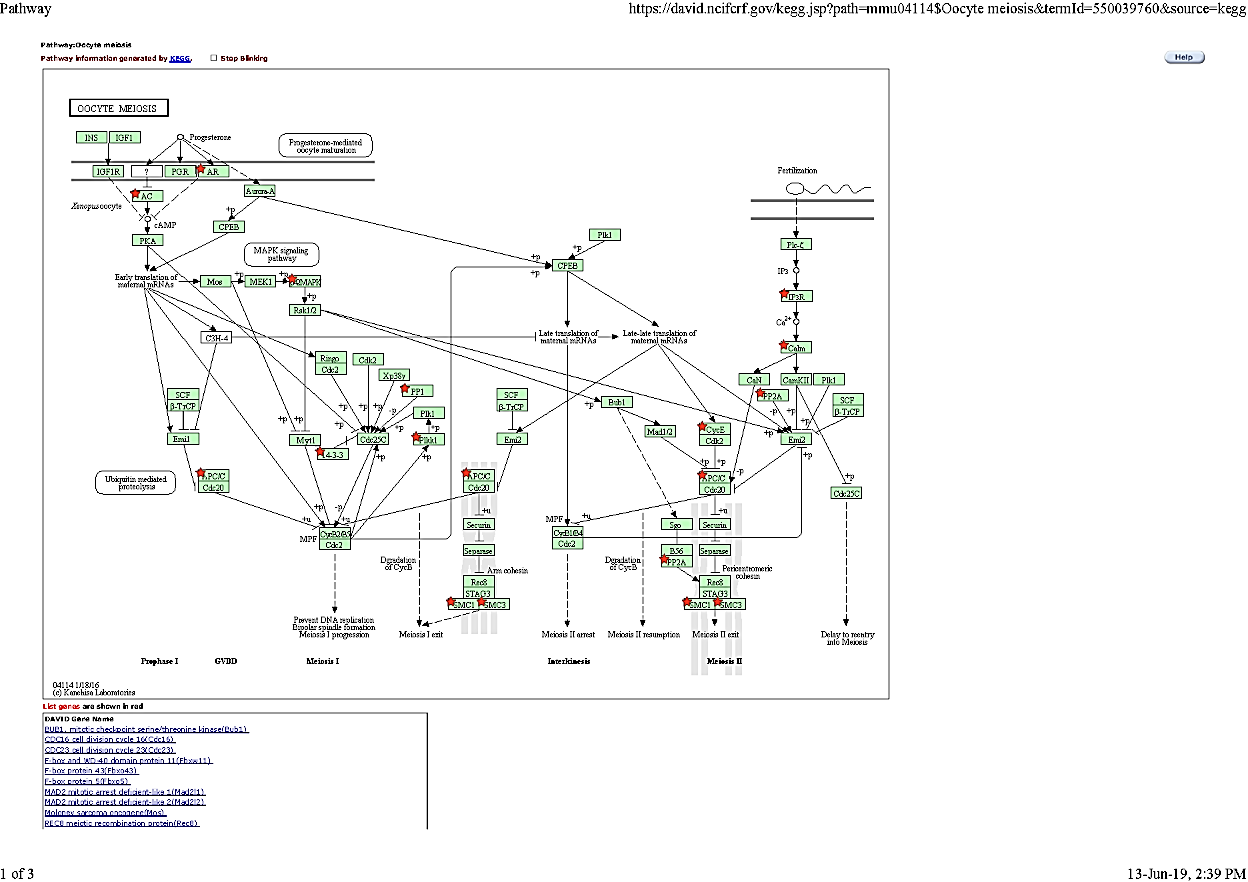

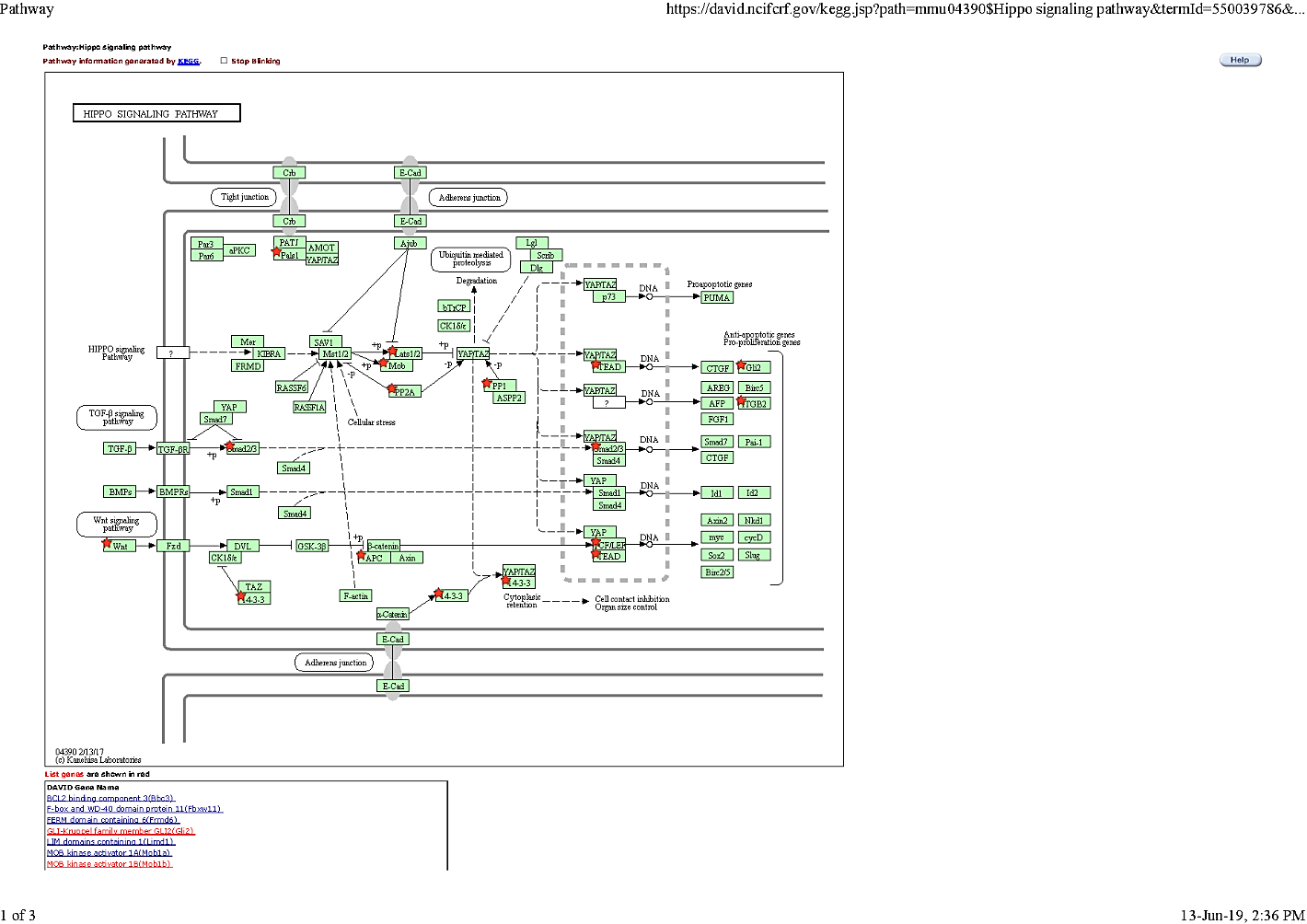

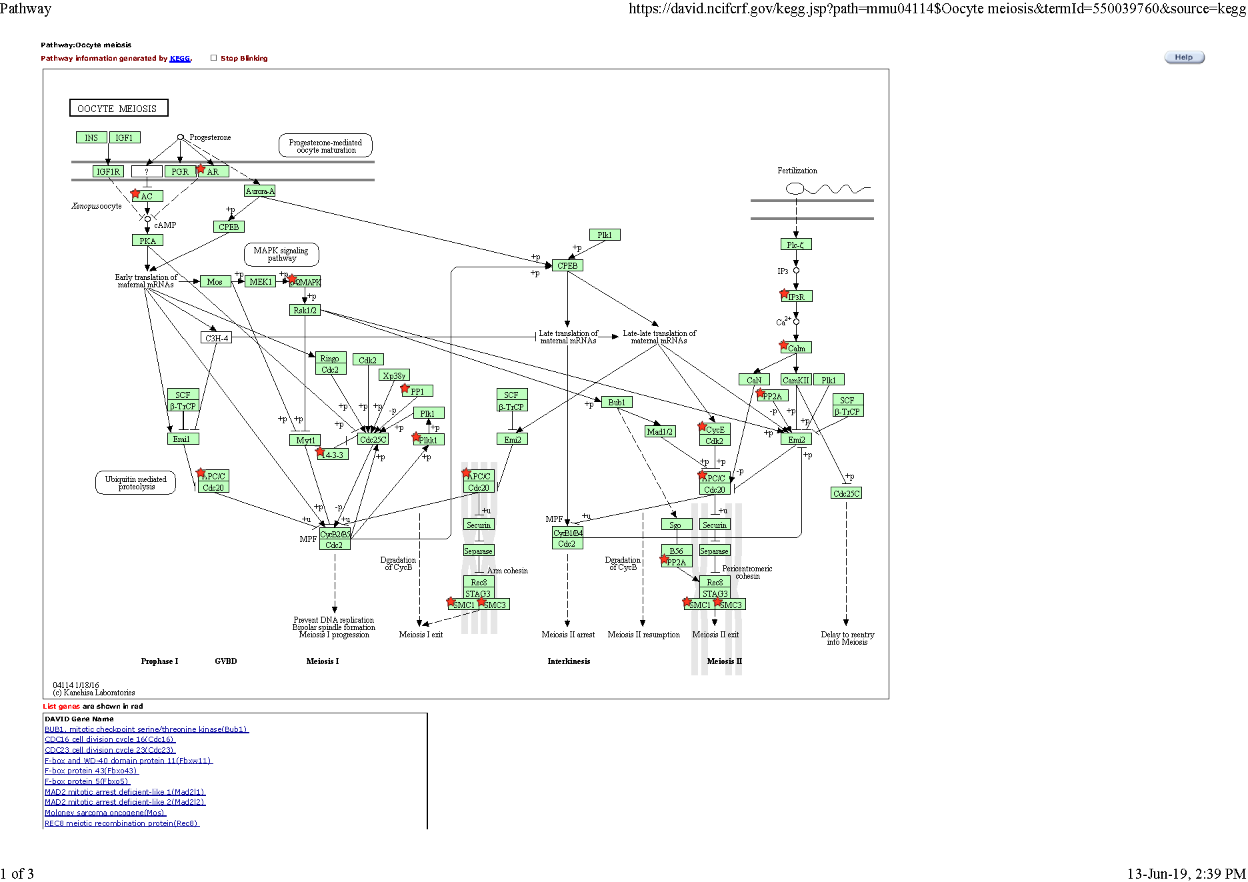

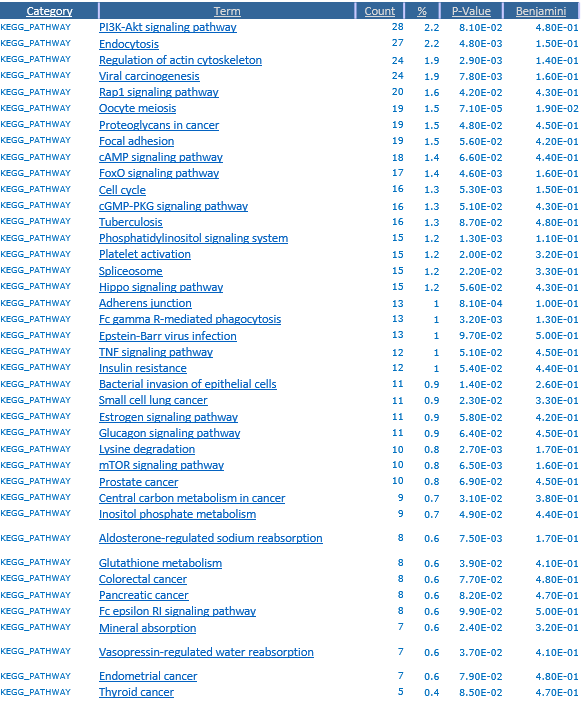

B

C

A

D

**Fig S2**: *Pathway analysis of differentially expressed genes in mice ovaries.* (A) Analysis was performed, using the Database for Annotation, Visualization and Integrated Discovery (DAVID), to determine the enriched pathways in the differentially expressed genes (DEGs) in the DSS-treated *vs* control mice, and the Kyoto Encyclopedia of Genes and Genomes (KEGG) pathways are listed. (B-D) Three of the most enriched pathways (B: PI3K-AKT; C: HIPPO; D: oocyte meiosis) are shown with the DEGs noted by red stars.

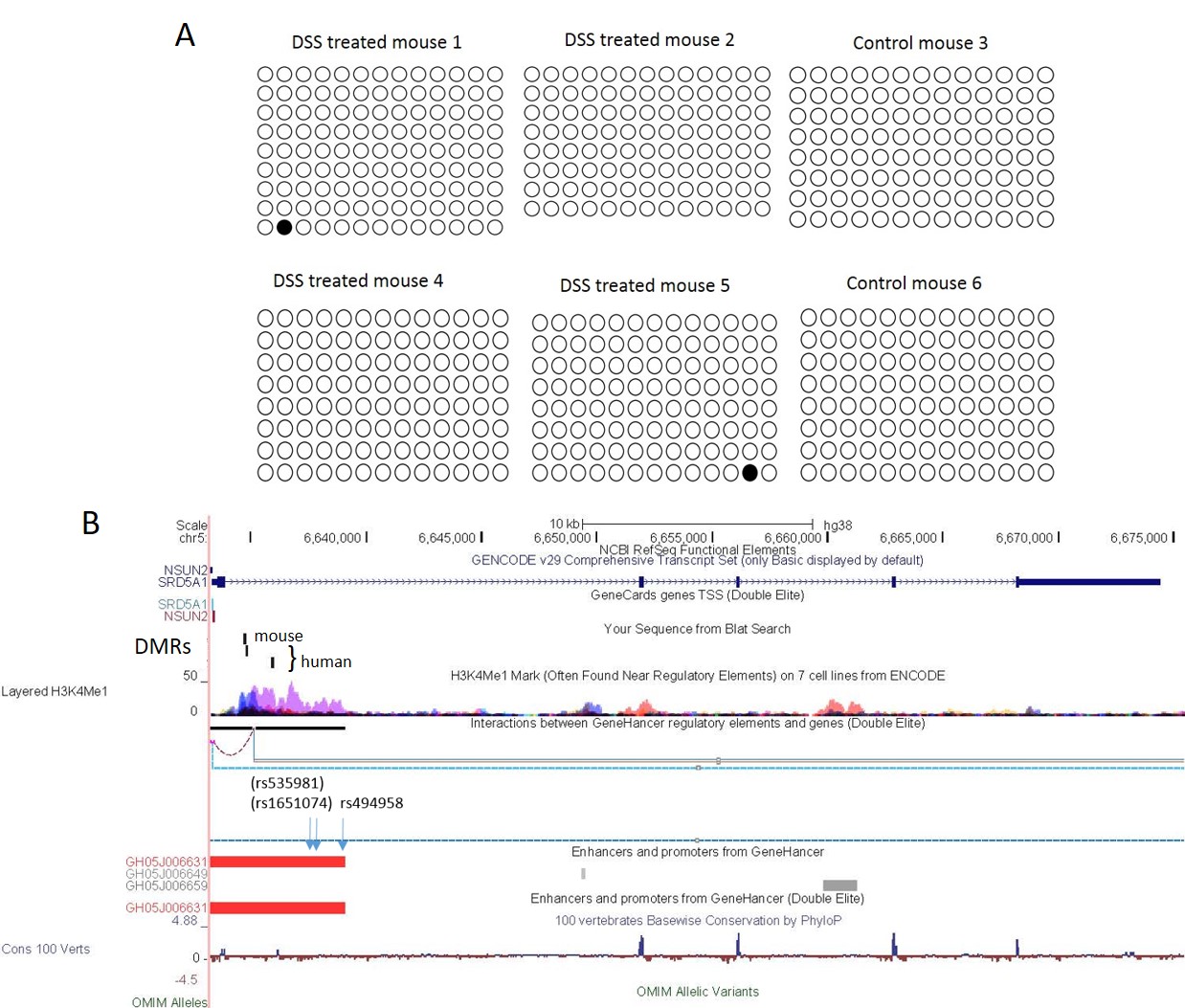

**Fig S3**: *Differential SRD5A1 methylation at a putative enhancer*. (A) After bisulfite conversion of ovarian DNA from DSS-treated (n=4) and control (n=2) mice, the *Srd5a1* proximal promoter was sequenced in multiple clones (rows). Each column represents one CpG (-123 to +62 relative to the TSS); white circles represent unmethylated CpGs and black ones are methylated. (B) The human *SRD5A1* genomic locus in the UCSC genome browser (GRC38/hg38) with observed DMRs marked, and homologous location of the DMR in the mice ovaries. Also shown are H3K4me1, GENEHancer elements, and an SNP (rs494958) associated with early menopause, and two others in high linkage disequilibrium.

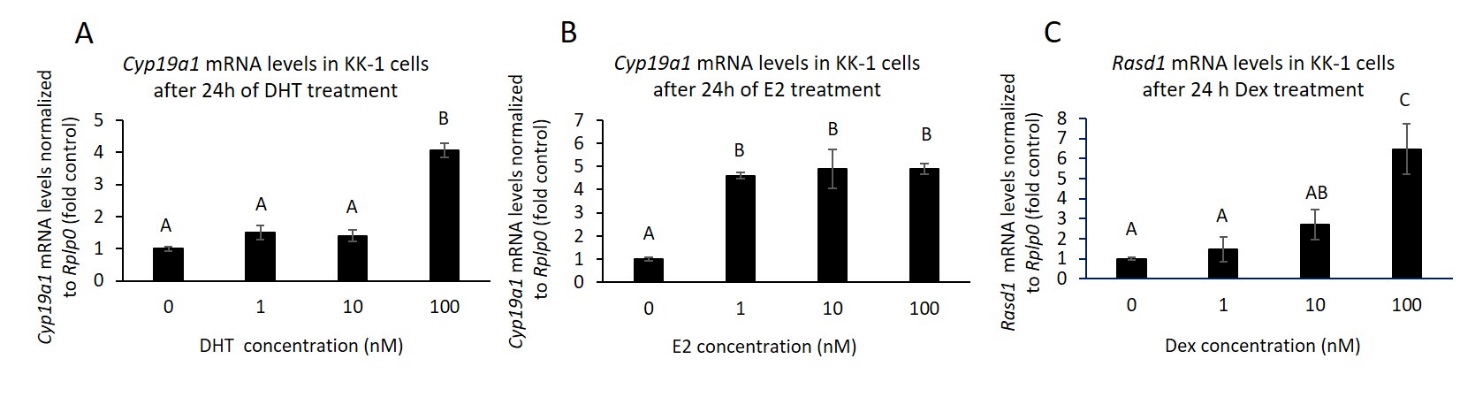

**Fig S4**: *Positive controls for steroid treatments in KK-1 murine ovarian cells.* (A-C) The mRNA from the experiments shown in Fig 4 of the main text was used to measure effects of the treatments on positive control genes (A,B) *Cyp19A1* and (C) *Rasd1*. Mean ± SEM is shown, with ANOVA followed by Tukey-Kramer t-test; means sharing the same letter are statistically similar (P>0.05).
